# Laboratory Methods Supporting Measles surveillance in Queensland, Australia, 2010 – 2017

**DOI:** 10.1101/472175

**Authors:** Jamie L. McMahon, Judy Northill, Mitchell Finger, Michael Lyon, Stephen B. Lambert, Ian M. Mackay

## Abstract

Australia was officially recognised as having eliminated endemic measles circulation in 2014. Maintaining laboratory support for surveillance of vaccine-preventable diseases such as measles is an essential component of reaching and maintaining circulation-free status. Between 2010 and 2017 over 13,700 specimens were tested in our laboratory by real-time RT-PCR (RT-rPCR). Positive extracts from travellers were sequenced as required. Sequences were uploaded to GenBank and demonstrated the wide genetic variation expected from the detection of MeV among virus introductions due to global travel. We describe the laboratory workflow employed in our laboratory between 2010 and 2017 for the sensitive detection of MeV infection, supporting high-quality surveillance.

## INTRODUCTION

The species *Measles morbillivirus* is an enveloped, single-stranded, non-segmented, negative-sense RNA member of the genus *Morbillivirus*, family *Paramyxoviridae*.^1^ Detection of measles virus (MeV) RNA by real-time RT-PCR (RT-rPCR) is a highly specific and sensitive rapid diagnostic tool to identify infection.^2-5^

Despite a safe and effective vaccine available for over 50 years, MeV remains a serious public health threat worldwide. There are hurdles in the path to global measles elimination and the eventual eradication of the MeV. In developing countries, a lack of political and financial support and areas of conflict give rise to low vaccine coverage through a limitation of access to adequate healthcare. In developed countries with little or no transmission, vaccine hesitancy and mistrust among parts of the population can cause gaps in immunity and, when concentrated in specific geographic locations, can result in local transmission and even sizable outbreaks after measles is introduced via infected travellers.^6^ Despite large gains in vaccination coverage resulting from the control initiatives, the global coverage for measles and rubella-containing vaccines has plateaued at 85% for the past ten years.^7^

Following MeV detection in the laboratory using RT-rPCR, genetic characterisation of MeV genotype primarily relies on sequencing a 450 nucleotide (nt) carboxyl-terminal region of the nucleocapsid (N450) protein and secondarily, on sequencing the entire coding region of the hemagglutinin genes, to produce internationally agreed upon measles genotype assignments.^8-10^

There is only a single serotype of MeV, but analysis of MeV genetic variation has identified eight MeV types (A–H) and 24 subtypes, referred to as genotypes.^9-11^ Not all genotypes are active. The World Health Organization (WHO) has declared six genotypes, not detected for at least 15 years, as officially inactive and there are a further five genotypes with no documented cases since 2006.^12^

RT-rPCR can also be designed to discriminate between wild-type MeV and vaccine-derived measles virus (MeVV). This approach is useful to distinguish between wild-type MeV infection and MeVV in individuals who received MeV-containing vaccine as post-exposure prophylaxis following exposure to wild-type virus. It may also be useful if a newly vaccinated person is infected by wild MeV very shortly after an initial routine vaccination was administered but before the development of a protective immune response that prevents future wild MeV infections.

We describe a workflow for the molecular diagnostic detection and characterisation of MeV in patients, particularly in Queensland, that has proven useful for supporting ongoing national efforts to maintain measles-free status in Australia.^13^

## METHODS

Samples for RT-rPCR testing were provided to this reference laboratory by hospitals and private laboratories from around the northeastern Australian State of Queensland. This testing, in response to clinical need, occurred as part of the laboratory’s role of supporting public health efforts to identify cases of measles, a vaccine-preventable disease. Detected strains were characterized by sequencing the N450 region as required. Throughout seven years, some earlier PCR reagents were replaced by newer, improved or preferred versions. The methods described below reflect the most recent reagents used. These changes did not affect the process.

### Nucleic acid extraction

Upon receipt of samples into the laboratory, viral RNA was extracted using the Qiagen EZ1 Mini kit v 2.0 (Qiagen, Hilden, Germany).

### Real-time RT-PCR screening and controls

Two RT-rPCRs were in use during the past decade to detect wild-type MeV and to discriminate MeVV from wild-type MeV. (Table 1 and Figure 1)

**Table 1.**
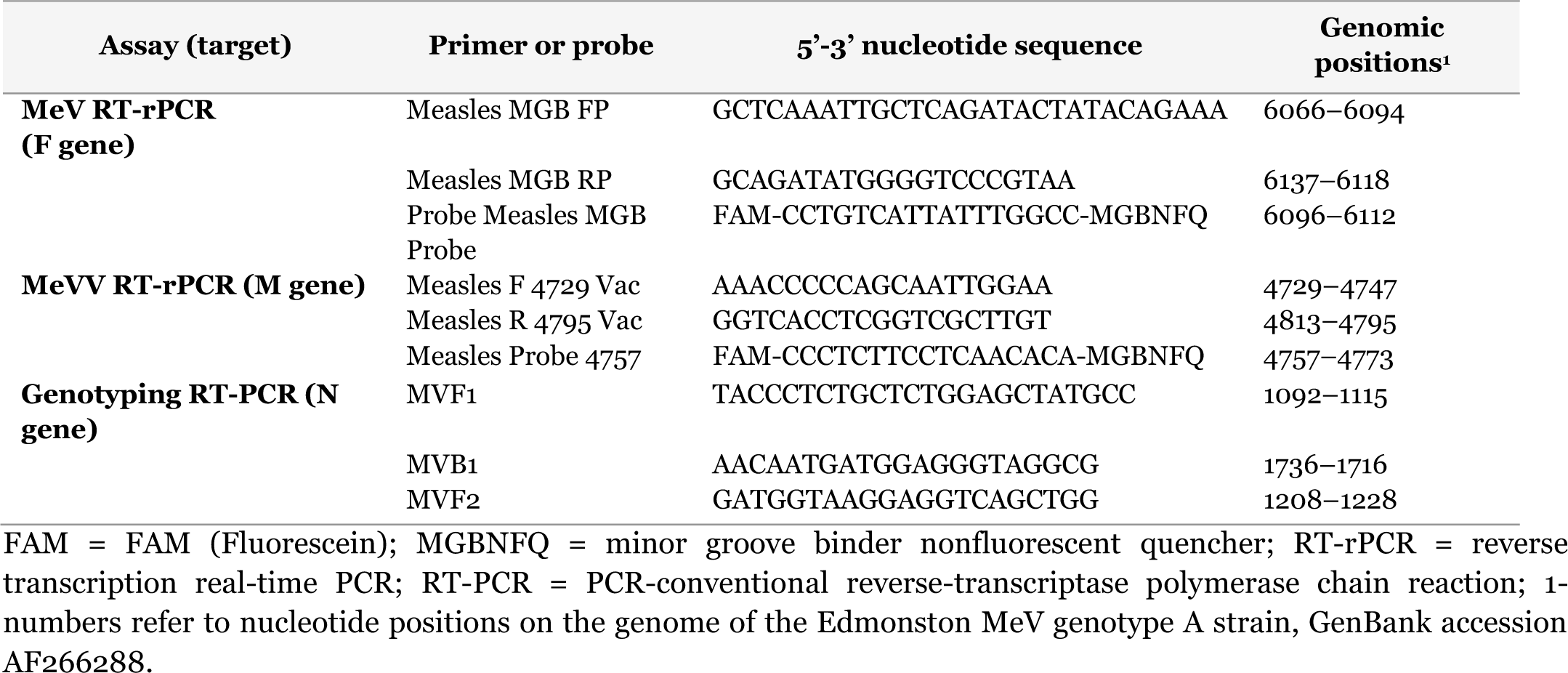
Oligonucleotides used to detect, differentiate and characterise MeV.

**Figure 1.**
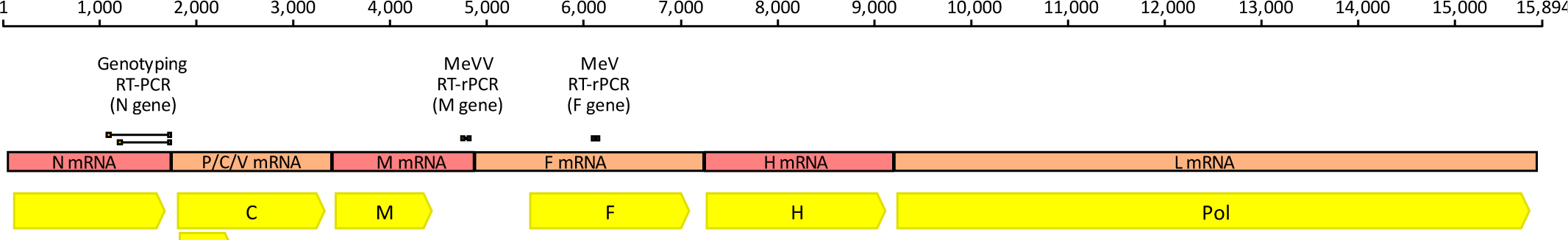
A generic MeV genome representation highlighting coding sequences, proteins produced (yellow boxes) and the diagnostic and sequencing PCR assay target regions. The assays indicated are described in Table 1. The genome is drawn to scale based on on MeV Edmonston strain (GenBank accession: AF266288). Higher resolution version is located at <u>10.6084/m9.figshare.7352468.</u>

The MeV RT-rPCR (Figure 1) was used as a screen for the detection of any *Measles morbillivirus*. The MeVV RT-rPCR was used as needed to discriminate circulating MeV genotypes from genotype A, which is not circulating at large but is used in MeV vaccines. However sequencing was the preferred option to confirm the presence of MeVV.

Synthetic primer and probe controls were developed using a method previously described.^14^ These were used for both RT-rtPCR assays, alongside no-template controls (NTC) and negative extraction controls. A threshold cycle (CT) of ≤40 was used to indicate detection of MeV RNA whereas CT values >40 were used to define a negative result.

### Conventional nested RT-PCR and sequence confirmation

Confirmation of selected RT-rPCR positive samples used a semi-nested conventional RT-PCR (N-gene RT-PCR) to generate a 450nt fragment encoding the carboxy-terminus of the nucleoprotein gene (N450).^15^ First-round RT-PC R amplification was performed after adding 5µl of RNA extract to 15µl SuperScript™ III One-Step RT-PCR System with Platinum™ Taq DNA Polymerase (Invitrogen, Carlsbad, CA) mixes. A 30 min reverse transcription incubation at 55°C and a 2 min inactivation at 94°C preceded 40 cycles of 95°C for 15 s, 60°C for 30 s, and 68°C for 60 s. A second round of amplification was performed by transferring 5µl of 1:100 diluted Round 1 product into 15µl Fast Cycling PCR Mix (QIAGEN, Hilden, Germany) reactions. A 5 min denaturation at 96°C preceded 40 cycles of 96°C for 5 s, 60°C for 5 s and 68°C for 18 s, with a final 72°C incubation for 60 s.

Initial N-gene amplicon analysis of second-round amplicon used traditional agarose gel electrophoresis in 0.5X TBE buffer, or the Qiagen QIAxcel (Qiagen, Hilden, Germany) according to the manufacturer’s instructions. For reactions producing the N450 product, gel electrophoresis of the remaining amplicon was followed by purification of the excised band using QIAquick Gel Extraction Kit (Qiagen, Hilden, Germany). Dye terminator sequencing was performed on the CEQ8000 Genetic Analysis System (SCIEX, Framingham, MA) or an ABI3130xl Genetic Analyser (Applied Biosystems, Australia) using the Big Dye terminator cycling ready reaction kit version 3.1 or GenomeLab DTCS Quick Start Kit (SCIEX, Framingham, MA).

### Subgenomic sequence analysis

Forward and reverse strands of raw sequence data were aligned with the WHO-designated MeV reference strains. Primer sequences were removed *in silico* using Sequencher software (various versions; Gene Codes Corporation). Sequence alignment for phylogenetic tree construction used Geneious version 8 and MEGA7 software, respectively.^16,17^

The phylogenetic tree was created using the Neighbor-Joining method with a bootstrap test of 500 replicates (Figure 2). Designation of the identification of the samples was confirmed using the MeaNS database genotyping tool.^8^

**Figure 2:**
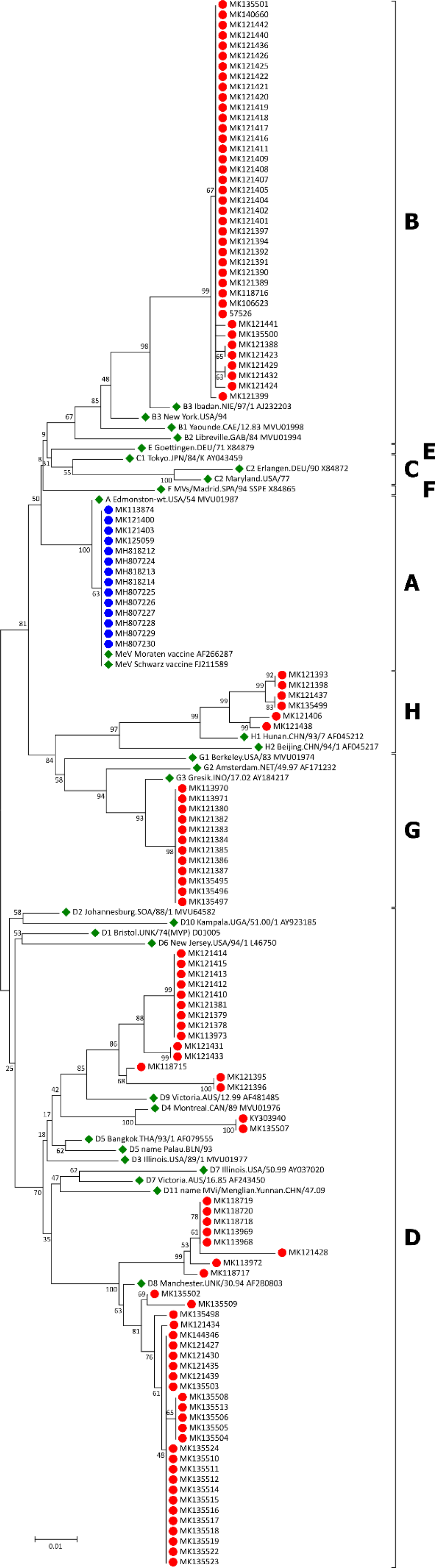
Phylogenetic analysis of partial nucleoprotein (N)-gene sequences from 122 MeV positive clinical specimens analysed in Queensland between 2010 and 2017. The sequenced region includes the World Health Organization-recommended N450 amplicon encoding the carboxyl-terminal of the N gene. MeV sequences associated with sequenced patients are marked in red, MeVV strain sequences in blue and WHO reference sequences, used to define genotypes and subtypes are marked in green.^12^ Sequences are labelled using GenBank accession numbers. The phylogenetic analysis used the Neighbor-Joining method in a bootstrap test (500 replicates) in MEGA7.^17,19-21^ A higher resolution version is located at <u>10.6084/m9.figshare.7352489</u>

### Ethical review

The Forensic and Scientific Services Human Ethics Committee assessed this project as not requiring full HREC review. It was not recognised to be research and is an audit of practice in accordance with the definition of research, page 6, of the National Statement on Ethical Conduct in Human Research (2007) updated in 2018).

## RESULTS

### MeV RT-rPCR (F gene) validation

The MeV RT-rPCR could reliably and repeatedly detect 20 copies per reaction determined after titrating a quantified *in vitro*-transcript and was linear across eight log10 dilutions of that transcript. The assay was precise (CT 35.0 with a standard deviation of 0.44 after amplifying 40 replicate reactions at the limit of detection).

The test detected 69 previously genotyped MeV positive samples including examples of clade A, B3, D2, D3, D4, D5, D8, G3, H1 and H2 MeV. The assay was 98% sensitive as it detected MeV in one sample that was IgG and IgM positive, but which could not be repeated for IgM and so was described as a false positive. The specificity of the test was evaluated by testing 65 MeV-negative samples including those known to be positive and negative for other viruses (enteroviruses, rubella virus and parvovirus) in nucleic acids extracted from a range of sample types (tissue culture, bone marrow, nasopharyngeal aspirates, nose swabs and throat swabs). The assay was 100% specific.

### MeVV RT-rPCR (M gene) validation

The MeVV RT-rPCR could reliably and repeatedly detect 240 copies per reaction determined after titrating a quantified *in vitro*-transcript and was linear across eight log_10_ dilutions of that transcript. The assay was precise after testing of repeatability (C_T_ 33.7 with a standard deviation of 1.6 after amplifying 33 replicate reactions near the limit of detection) and reproducibility (C_T_ 33.1 with a with a standard deviation of 1.9 after amplifying 28 reactions near the limit of detection on different days, by different operators using different reaction mix batches.

The test detected eight of eight previously genotyped MeV clade A positive samples. The assay was 100% sensitive. In additional specificity testing, the MeVV test was evaluated by testing 92 MeV clade A-negative samples. These included extracts known to be positive and negative for MeV non-clade A viruses (from among 35 genotyped MeV examples of clade A, B3, D2, D3, D4, D5, D8, G3, H1 and H2 MeV) and for entirely different viral species (enteroviruses, rubella virus and parvovirus) in nucleic acids extracted from a range of sample types (tissue culture, bone marrow, nasopharyngeal aspirates, nose swabs and throat swabs). The assay was 100% specific.

### Summary of use between 2010-17

Between 2010 and 2017, we received 13,718 samples for MeV RT-rPCR testing.

Most samples were swabs (71.0%), urine (25.5%) and aspirates (2.7%). Cerebrospinal fluid, blood and washings comprised a small proportion (0.8%). Of these, 533 samples tested positive by MeV RT-rPCR. The MeVV RT-PCR, which was used sporadically from 2012 onwards, identified 141 positive detections from 127 cases (5 positive samples from 5 cases in 2012, 36 from 33 cases in 2013, 42 from 38 cases in 2014, 21 from 19 cases in 2015 24 from 22 cases in 2016 and 15 from 15 cases in 2017). Most measles cases in Australia are due to direct importation, or contact with travel-related cases.^13^

Samples in this study were collected in Queensland, from travellers who had returned to Queensland from international sites (the Solomon Islands, Papua New Guinea, Indonesia, India, China, Myanmar, Vietnam, the Philippines Thailand, Pakistan), or who visited Queensland from interstate (New South Wales, our neighbouring state). Contacts of travellers and cases were also tested as required.

After accounting for multiple sample types tested per person, 340 positive RT-rPCRs remained. Genotyping was not performed on every positive sample. Some were part of clusters where the genotype in contacts was expected to be identical to that from the initial case. Some MeV positives were not subjected to sequencing as they were expected to represent virus shed due to a contemporaneous vaccination, and thus were likely to be MeV genotype A. One hundred and twenty-two MeV strains were characterised by nucleotide sequencing, and the N450 sequences submitted to GenBank and analysed alongside prototype genotype sequences (Figure 2). Among the sequences were confirmed cases of genotype A, indicating detection of MeVV strain (indicated in blue; Figure 2) post-vaccination. Some of these were included in submissions to GenBank but their N450 sequences were unremarkable.

Apart from genotype A MeVV strain, N450 sequencing between 2007 and 2010 identified MeV genotypes B3, D4, D8, D9, G3 and H1, like those described by other reports during similar periods.^9,13,18^ No strains of genotype B1-2, C1-2, D1-3, D5-7, D10-11, E, F, G1-2 or H2 were identified. We did not detect any genotypes considered extinct (B1, C1, D1, E, F, G1)^12^ or inactive (D2, D3, D10, G2, H3).^10^

## CONCLUSION

Australia was officially recognised for eliminating endemic measles transmission in 2014 by providing evidence of no endemic circulation of measles virus for at least three years before this.^22^

We have described a laboratory workflow employed in Queensland, Australia, across this time point, to detect imported cases by supporting high-quality surveillance. This workflow successfully detected and genotyped MeV infections in Queensland between 2010 and 2017, in response to clinical need. A low proportion of all samples tested were positive, and of those, a portion was sequenced. Among the sequenced MeV strains were a wide range of genotypes as expected in a setting where endemic measles transmission has been halted. This genetic diversity reflects MeV introductions originating from a range of international travel sources that have yet to eliminate measles. from a range of international travel sources that have yet to eliminate measles.

A low proportion of all samples tested were positive, and of those, a portion was sequenced. Among the sequenced MeV strains were a wide range of genotypes as expected in a setting where endemic measles transmission has been halted. This genetic diversity reflects MeV introductions originating from a range of international travel sources that have yet to eliminate measles.

Ongoing MeV mutation can challenge molecular diagnostic assays, resulting in reduced test performance or even test failure because of mismatches between oligonucleotides and target. We recently updated our MeV RT-rPCR test to better accommodate viral evolution. Optimisation and validation of the improved RT-rPCR test is ongoing and will be described elsewhere.

Most of our MeV positives were the result of an infection acquired during travel to Australia from countries with ongoing endemic measles transmission: either from returned Australian travellers and their contacts or from specimens provided for testing/confirmation directly from these countries as part of our reference laboratory service. High-quality laboratory services are required in settings where endemic measles circulation has been halted to identify imported virus and its transmission and to confirm continuing elimination status.

## ACKNOWLEDGEMENTS

We thank Public Health Virology members Russell Simmons, Frederick Moore, Glen Hewitson, Doris Genge, and Peter Burtonclay who contributed to the laboratory testing of the cases. We also thank the staff in the public health units involved in following up the cases and their families.

## REFERENCES

1. Wang L.-F., Collins P.L. Fouchier, R.A.M., Kurath G., Lamb R.A., Randall R.E., Rima B.K. Paramyxoviridae. ICTV 9th Report (2011) https://talk.ictvonline.org/ictv-reports/ictv_9th_report/negative-sense-rna-viruses-2011/w/negrna_viruses/199/paramyxoviridae. Accessed 11/11/2018, 2018.

2. Benamar T., Tajounte L., Alla A., et al. Real-Time PCR for Measles Virus Detection on Clinical Specimens with Negative IgM Result in Morocco. PLoS One. 2016;11(1):e0147154.

3. Thomas B., Beard S., Jin L., Brown K. E., Brown D. W. Development and evaluation of a real-time PCR assay for rapid identification and semi-quantitation of measles virus. J Med Virol. 2007;79(10):1587–1592.

4. Sanz J. C., Mosquera M., Ramos B., Ramirez R., de Ory F., Echevarria J. E. Assessment of RNA amplification by multiplex RT-PCR and IgM detection by indirect and capture ELISAs for the diagnosis of measles and rubella. APMIS. 2010;118(3):203–209.

5. Michel Y., Saloum K., Tournier C., et al. Rapid molecular diagnosis of measles virus infection in an epidemic setting. J Med Virol. 2013;85(4):723–730.

6. Lo N. C., Hotez P. J. Public Health and Economic Consequences of Vaccine Hesitancy for Measles in the United States. JAMA Pediatr. 2017;171(9):887–892.

7. Moss W. J. Measles. Lancet. 2017;390(10111):2490–2502.

8. Rota P. A., Brown K., Mankertz A., et al. Global distribution of measles genotypes and measles molecular epidemiology. J Infect Dis. 2011;204 Suppl 1:S514–523.

9. World Health Organization. The role of extended and whole genome sequencing for tracking transmission of measles and rubella viruses: report from the Global Measles and Rubella Laboratory Network meeting, 2017. Wkly Epidemiol Rec. 2018;93(6):55–59.

10. World Health Organization. Measles virus nomenclature update: 2012. Wkly Epidemiol Rec. 2012;87(9):73–81.

11. Rota P. A., Featherstone D., Bellini D.J. Molecular Epidemiology of Measles Virus. In: Griffin DE, Oldstone MM, eds. Measles. Pathogenesis and control. Introduction. Vol 330. 2009/02/11 ed. Berlin Heidelberg: Springer-Verlag; 2009.

12. Munoz-Alia M. A., Muller C. P., Russell S. J. Antigenic Drift Defines a New D4 Subgenotype of Measles Virus. J Virol. 2017;91(11).

13. Gidding H. F., Martin N. V., Stambos V., et al. Verification of measles elimination in Australia: Application of World Health Organization regional guidelines. J Epidemiol Glob Health. 2016;6(3):197–209.

14. Smith G., Smith I., Harrower B., Warrilow D., Bletchly C. A simple method for preparing synthetic controls for conventional and real-time PCR for the identification of endemic and exotic disease agents. J Virol Methods. 2006;135(2):229–234.

15. Chibo D., Birch C. J., Rota P. A., Catton M. G. Molecular characterization of measles viruses isolated in Victoria, Australia, between 1973 and 1998. J Gen Virol. 2000;81(Pt 10):2511–2518.

16. Kearse M., Moir R., Wilson A., et al. Geneious Basic: an integrated and extendable desktop software platform for the organization and analysis of sequence data. Bioinformatics. 2012;28(12):1647–1649.

17. Kumar S., Stecher G., Tamura K. MEGA7: Molecular Evolutionary Genetics Analysis Version 7.0 for Bigger Datasets. Mol Biol Evol. 2016;33(7):1870–1874.

18. Woudenberg T., van Binnendijk R. S., Sanders E. A., et al. Large measles epidemic in the Netherlands, May 2013 to March 2014: changing epidemiology. Euro Surveill. 2017;22(3):pii: 30443.

19. Saitou N., Nei M. The neighbor-joining method: A new method for reconstructing phylogenetic trees. Mol Biol Evol 1987;4(4):406–425.

20. Tamura K., Nei M., Kumar S. Prospects for inferring very large phylogenies by using the neighbor-joining method. Proc Natl Acad Sci USA. 2004;101(30):11030–11035.

21. Kumar S., Tamura K., Nei M. MEGA3: Integrated software for molecular evolutionary genetic analysis and sequence alignment. Briefings in bioinformatics. 2004;5(2):150–163.

22. World Health Organization Regional Office for the Western Pacific. Third Annual Meeting of the Regional Verification Commission for Measles Elimination in the Western Pacific, Seoul, Republic of Korea, 18–21 March 2014: report. 2014. http://iris.wpro.who.int/handle/10665.1/10675. Accessed 14/11/2018.

